# Cross-Kingdom Genomic Conservation of Human Sleep-Related Gene Orthologs: Phylogenomic Evidence from *Chlamydomonas reinhardtii*

**DOI:** 10.1101/2024.11.02.621661

**Authors:** Seithikurippu R Pandi-Perumal, Konda Mani Saravanan, Sayan Paul, George C. Abraham, David Warren Spence, Saravana Babu Chidambaram

## Abstract

Sleep is a widespread and evolutionarily conserved process observed in diverse organisms, from jellyfish to mammals, hinting at its origin as a life-supporting mechanism over 500 million years ago. Although its fundamental purpose and mechanisms remain unclear, the evolution and adaptive significance of sleep continue to be debated. This study explores the evolutionary origins of sleep using Chlamydomonas reinhardtii as a model organism, identifying 145 orthologs analogous to known sleep-related genes across species, highlighting the evolutionary conservation of sleep-regulating pathways. Additionally, discovering uncharacterized proteins with high sequence similarity and significant e-values suggests unexplored roles in sleep regulation, underscoring the potential of C. reinhardtii to reveal new insights into the molecular basis of sleep. This work provides a foundation for identifying previously unknown sleep-associated proteins, particularly within single-celled organisms, which may offer novel perspectives on the biological role of sleep. The study demonstrates that phylogenomic analysis of diverse model organisms can expand our understanding of the evolutionary trajectory of sleep and its fundamental function, paving the way for further research in sleep biology and its health implications. Overall, the fundamental functions of sleep observed in higher animal phyla originated from its primordial activities, demonstrating an evolutionary continuum wherein more specialized tasks were integrated with sleep’s essential restorative properties.

---

The most fundamental and essential inquiries regarding sleep are: Why do we sleep? Is sleep a vital necessity? Do all living things sleep? Are hibernation, sleep, and dormancy tactics exclusive to higher phyla, particularly mammals and seed plants? Are these strategies that have helped adapt to combat the adversities of climate? The question, “Should we view sleep as a characteristic of entrained rhythm?” provides a starting point around which one can discuss the significance of sleep as a vital component of life that permits living organisms to survive on planet Earth. It is undisputed that organisms get re-stimulated to act, rejuvenate, and prolong their stay on earth due to the recharging effect of sleep. This evolutionary perspective on life supports the notion that sleep has advanced as a trait because it has perhaps facilitated the process of life colonizing the land.

Occurring almost universally in land-dwelling forms of life, sleep forms a part of the endogenous rhythm of living beings, all indispensably guided by the metronome of this planet’s repetitive cycles, manifesting as entrained behavior. Plants and animals get accustomed to this time-locked behavior subjected to diurnal stimuli and environmental cues. The sleep mechanism draws a connation with photo and skotomorphogenesis. Investigating the reasons behind sleep determines its essentiality and calls for a holistic understanding.

Deciphering the reason that sleep appears to be an evolutionarily conserved trait, thus suggesting that it is essential for survival, has been a major puzzle for “sleep” scientists. Sleep of a vitally important trait, and one can persuasively argue that it serves some important needs, although scientists have still not worked out what this need is. This might prompt the suggestion that if scientists have failed to detect the purpose of sleep, then perhaps it doesn’t have a purpose. Not everyone agrees with this nihilistic view. As Allan Rechtschaffen pointed out, “*if sleep does not serve an absolutely vital function, then it is the biggest mistake the evolutionary process has ever made*” (Rechtschaffen Allan 1971). One possible function that sleep may serve is to restore and rejuvenate the organism. This is considered the “energy conservation” view (Berger and Phillips 1995; Latifi et al. 2018). This strategy facilitates efficient energy distribution, promoting sustained restful sleep (Siegel 2005). Nevertheless, it could be argued that the strategy of reducing energy consumption also increases the vulnerability to environmental dangers in as much as perceptual vigilance produces a significant drain on energy stores. This risk therefore undercuts the argument that sleep acts to promote survival. The general conceptualization that sleep is a vital, life-supporting function has therefore not received universal support among scientists.

The skepticism of the anti-conservation view of sleep is further supported by the observation that non-sleeping fish do achieve sleep’s major benefits by somehow circumventing the necessity for sleep altogether (Kavanau 2008). These perspectives and their associated tools have no restrictions concerning the species they study and thus can investigate the full gamut of the living world, including the lowest of forms of life. One such species, with this particular interest, is the type of material of the current investigation. *Chlamydomonas reinhardtii,* the focal organism of study, is a tiny phytoplankton. This unicellular eukaryotic volvocine, often known as a chlorophycean, is a single-celled green alga etymologically named Chlamydomonas (Greek *Chlamys*, a cloak or mantle; *monas*, solitary). It is recognized as a progenitor from which a lineage of terrestrial plants split off more than a billion years ago (Merchant et al. 2007).

Chlamydomonas is widely regarded as a valuable model organism in the fields of genetics and molecular biology for several reasons. It provides a wealth of information by serving as a genomic resource with wide collections of knockout strains offering scope for CRISPR (clustered regularly interspaced short palindromic repeats) gene editing techniques (Bharathkumar et al. 2021). These resources are characterized by their simplicity (simple life cycle), speed (easy isolation of mutants), accuracy (owing to the single-celled state with multiple affiliations connecting plant, animal, and microbial world), efficiency (expanding amenability to multiple sets of tools and techniques), cost-effectiveness (attributed to photoautotrophic survival in culture), and the versatility (in approach and handling to address various challenging questions) allow this organism to provide a wide range of research options (Harris 2001). These advantages support claims that *Chlamydomonas reinhardtii* (herein referred to as Chlamydomonas) is the most convenient and versatile microbial platform for the investigation of basic physiological and behavioral processes (Blaby et al. 2014).

The genome of *C. reinhardtii* has been extensively studied due to its significance in elucidating evolution, photosynthesis, and biotechnology. The Chlamydomonas genome draft (v3) released in 2007 (Merchant et al. 2007) confirmed that the nuclear DNA uncovers over 99 percent of all *C. reinhardtii* genes. Approximately 19,500 proteins are encoded by the nuclear genome, sized in about 120 megabytes on 17,741 loci with 19,526 putative transcripts. It accounts for a variety of roles in processes such as base metabolism, flagellum construction, and photosynthesis. Additionally, Chlamydomonas contains DNA from mitochondria (chondriome) and chloroplasts (plastome), and serves in comprehending integrated functions, more specifically in deciphering the genetic and functional makeup of the two organelles. Chlamydomonas is a genus of single-celled biflagellate green microalgae, ironically called a ’green yeast’ or ’photosynthetic yeast’, due to its semblance with *Saccharomyces cerevisiae* (Rochaix 1995; Mittag et al. 2005). It stands out as the most thoroughly studied and the most prominent model organism in a variety of scientific domains. Chlamydomonas is often referenced in the scientific literature for its potential in biofuel production and is consequently widely examined in other commercially viable processes (Scranton et al. 2015). Its relevance to sleep studies is that its flagellae contains an eye spot that exhibits a pronounced circadian rhythm (BRUCE 1970; Mittag et al. 2005; Matsuo et al. 2008; Matsuo and Ishiura 2010; Niwa et al. 2013). With the help of the eyespot apparatus, Chlamydomonas can swim toward or away from the light source and find a favorable environmental niche to maximize its photosynthetic development (Yoshimura and Kamiya 2001).

Since Dangeard’s cytological findings of haploid chromosome count in 1899, Chlamydomonas has been a popular organism for studies on genomic regulation and interactions (Levine and Ebersold 1960). Over the years, the species *C. reinhardtii* has been proven to be a valuable model organism with many advantages because of its rapid axenic development despite its capacity to grow in the dark, resemblance to mammals in terms of protein structure, metabolic profile, and its constituents with specifics known in terms of its gene content and physiological traits, and ease of genetic manipulation (Sasso et al. 2018; Findinier and Grossman 2023).

Information on nuclear (Merchant et al. 2007), chloroplast (Maul et al. 2002), and mitochondrial DNA (Vahrenholz et al. 1993) have been published and documented quite well. Using documented data as a starting point, this present inquiry focuses on sleep genes whose presence is likely to have relevance to sleep research.

## METHODS

### COLLECTION OF SLEEP GENES

Genes associated with sleep, sleep function, and sleep disorders were manually gathered from the NCBI database. The pertinent keywords identified, used either alone or in combination with words or phrases (such as sleep, circadian rhythm, insomnia, sleep disorder, sleep regulation, and sleep-related genes) helped generate a host of data that were scrutinized for broader relevance sleep research. Sequence searches were performed using the listed specific keywords, and the outcome was narrowed down by utilizing filters such as ‘Homo sapiens’ to concentrate on genes related to humans, as well as other pertinent filters such as ‘publication date ranges’ and ‘research type’ to keep the study consistent with relevant research goals. Gene summaries examined and associated literature for each outcome, with particular attention to annotations, helped to indicate directed one-to-one participation with sleep processes or sleep disorders. An exhaustive compilation of identified genes was created, as well as documentation of their symbols, complete names, and functional annotations. The involvement of these elements in sleep function was verified by cross-referencing the list with scholarly publications using NCBI’s PubMed database (Jin et al. 2024). Supplementary sources such as GeneCards and Online Mendelian Inheritance in Man (OMIM) were used to verify the genes more thoroughly and compile new sleep research reviews (Hamosh et al. 2021). A carefully selected compilation of genes obtained by following this methodological approach alluded to the unraveling of roles played in sleep genes. It is expected that the outcome might serve as a basis for follow-on studies and analyses.

#### Blast Analysis of Sleep Genes against *Chlamydomonas reinhardtii genome*

BLAST analysis of sleep genes against the *C. reinhardtii* genome was carried out. Through BLAST analysis of sleep genes (Altschul et al. 1990), homologous sequences in the *C. reinhardtii* genome are found, demonstrating the evolutionary relationships between *C. reinhardtii* and other species. The nucleotide or protein sequences of the discovered sleep-related genes from the NCBI database were collected, with vigilance for their consistency with the required FASTA format. The NCBI BLAST homepage (URL: https://blast.ncbi.nlm.nih.gov/Blast.cgi) was accessed, followed by the use of suitable BLAST software based on the type of analyzed sequences, such as BLASTn for nucleotide sequences or BLASTp for protein sequences. On specifying the desired database by opting for *C. reinhardtii* genome data from NCBI, to the extent it is accessible, or by acquiring it from alternative sources and utilizing the ‘align two or more sequences’ feature.

Sleep gene sequences in the BLAST interface were obtained by uploading or pasting them, which also formed part of the working strategy. Following this, the search parameters were adjusted by including the E-value threshold, match/mismatch scores, gap penalties, and word size. A BLAST search was performed to provide a basis for analyzing the results, specifically paying attention to alignment scores, E-values, and percent identity to discover essential matches. On examining the biological importance of the detected similarities were assessed by comparing them with values in the current literature to functional consequences of the homologous genes in *C. reinhardtii*, and from this, the base material was created. Eventually, the procured data was collated and recorded with notes on discoveries. Encompassing results on BLAST parameters, a concise overview of results, and an explanation accompanied by visual depictions such as alignment graphs substantiated the investigation. This systematic approach provided highly useful insights into the evolutionary preservation and functional resemblances of genes associated with sleep across different species.

### ORTHOLOGY AND FUNCTIONAL ANNOTATION (EGGNOG) OF SLEEP GENES

Based on orthology and functional annotation by EggNOG analysis, sleep-related genes are characterized according to their evolutionary relationships and functions (Huerta-Cepas et al. 2017). In conducting evolutionary genealogy of genes, the Non-supervised Orthologous Groups (EggNOG) annotation of sleep genes and the nucleotide or protein sequences of the selected sleep-related genes from the NCBI database verified in the FASTA format were retrieved. This was accomplished by utilizing the site of the EggNOG database (URL: http://eggnog5.embl.de) and the EggNOG-mapper program (Jensen et al. 2008). The sleep gene sequences were directly pasted into the input box provided on the EggNOG-mapper interface (Cantalapiedra et al. 2021). Search parameters were customized by choosing the suitable taxonomic scope and adjusting to settings such as E-value thresholds. Thus, the EggNOG-mapper program was utilized to perform sequence mapping to EggNOG orthologous groups, and the functional annotations were applied based on conserved domains and known functional information.

The results were examined for functional descriptions in Gene Ontology (GO) terms, KEGG pathways, and COG categories linked to each sleep gene (Kanehisa and Goto 2000; Tatusov et al. 2003). By stressing conserved domains and pathways, functional annotations as sleep genes were examined. The functional roles of genes involved in sleep processes were validated by cross-referencing the EggNOG annotations with current literature. A thorough report on the detailed overview of the EggNOG annotation results was compiled. This report included the gene sequences specific to EggNOG-mapper settings with a concise explanation of the functional annotations. Additionally, pathway diagrams were utilized for visual representation and the distributions of GO terms. The application of this systematic approach may provide insight into how sleep-related genes ultimately regulate biological functions.

### INTERPRO SCAN OF SLEEP GENES

A focused proteomic study of sleep genes reveals conserved domains, protein families, and functional indicators implicated in sleep-regulated biological processes. In conducting an InterProScan analysis on sleep genes, the nucleotide or protein sequences of the identified sleep-related genes were obtained from the NCBI database and their placement in the appropriate FASTA format was verified. The InterProScan tool on the EMBL-EBI website (https://www.ebi.ac.uk/interpro/interproscan.html) was accessed and chosen as a suitable tool for analyzing protein sequences to detect domains, namely selected families and functional sites (Quevillon et al. 2005). The sleep gene sequences in FASTA format were submitted by uploading the file and copying and pasting the sequences into the designated input box on the InterProScan interface. The search parameters were customized by picking the desired types of analysis to execute (such as Pfam, PRINTS, and PROSITE patterns)(Attwood 2002; Hulo et al. 2006; Mistry et al. 2021), fine-tuning the sensitivity, and selecting the preferred output formats (such as XML, TSV, and HTML). The InterProScan program was utilized to analyze the input sequences. These analyses involved the identification of functional domains, signatures, and protein families by integrating prediction models from several databases. Specific attention was paid to the intricate details regarding protein domains, families, functional sites, and other pertinent annotations linked to each sleep gene. Functional annotations were examined to deepen the understanding of the functions of sleep genes, with a focus on conserved domains, active sites, and other recognized functional characteristics. The functional aspects of InterProScan annotations were validated by cross-referencing them with contemporary data.

This technique catered to gain unique insights into the genome involvement and regulation in sleep processes. Combined with due notes on the gene sequences, specific InterProScan parameters were utilized to arrive at an objective explanation of the functional annotations. Additionally, visual representations such as domain architectures and functional sites to enhance the understanding of the results were incorporated. This line of inquiry offered a comprehensive functional annotation and added to the knowledge of the biological functions of sleep-associated genes.

### PATHWAY ANALYSIS OF SLEEP GENES

Sleep gene pathway analysis identifies molecular networks and biological pathways essential for sleep regulation and associated processes. Sleep-related genes and their identifiers (such as gene symbols or Entrez IDs) were compiled to analyze sleep genes’ pathways. Pathway analysis tools, such as KEGG, Reactome, Ingenuity Pathway Analysis (IPA), or DAVID were selected as the chosen as instruments of this study (Kanehisa and Goto 2000; Dennis et al. 2003; Croft et al. 2011; Krämer et al. 2014). The list of genes relevant to the investigation was entered to ensure compatibility with the format. Then, the analysis settings were adjusted by selecting the organism of interest and the desired type of analysis using pathway enrichment and network analysis. Pathway analysis was performed to associate the genes with established biological pathways and detect pathways that show significant enrichment. The result was analyzed by specifically examining the enriched pathways with their implicated genes and associated statistical significance. The biological importance of the discovered pathways was examined, and their relevance to sleep processes was assessed. Validation of findings was carried out by cross-referencing them with the existing literature. To cap the study by generating a comprehensive report on the pathway analysis, information about input genes, analysis parameters, enriched pathways, and their statistical significance were determined. Visual representations such as pathway diagrams and network maps were included.

## Results

### Orthologs of Sleep Genes in Chlamydomonas

Table 1 displays the results of the BLAST analysis performed to identify the existence of orthologous genes in *C. reinhardtii* associated with sleep. The acquired findings encompassed identified and unidentified proteins, potentially providing insights into preserving sleep pathways across diverse species. An observation that can be made from the table is that there is a significant proportion of proteins that are yet to be characterized, as shown by the CHLRE IDs. Although not documented in the literature, these proteins exhibit homology to the sleep-related genes based on the e-values and the sim mean score. An illustrative example is the protein CHLRE_16g686650v5, currently uncharacterized but has a significant e-value of 4. The first measurement is 21E-51, indicating a minimal value that reflects the similarity of 50.58%.

**Table 1.**
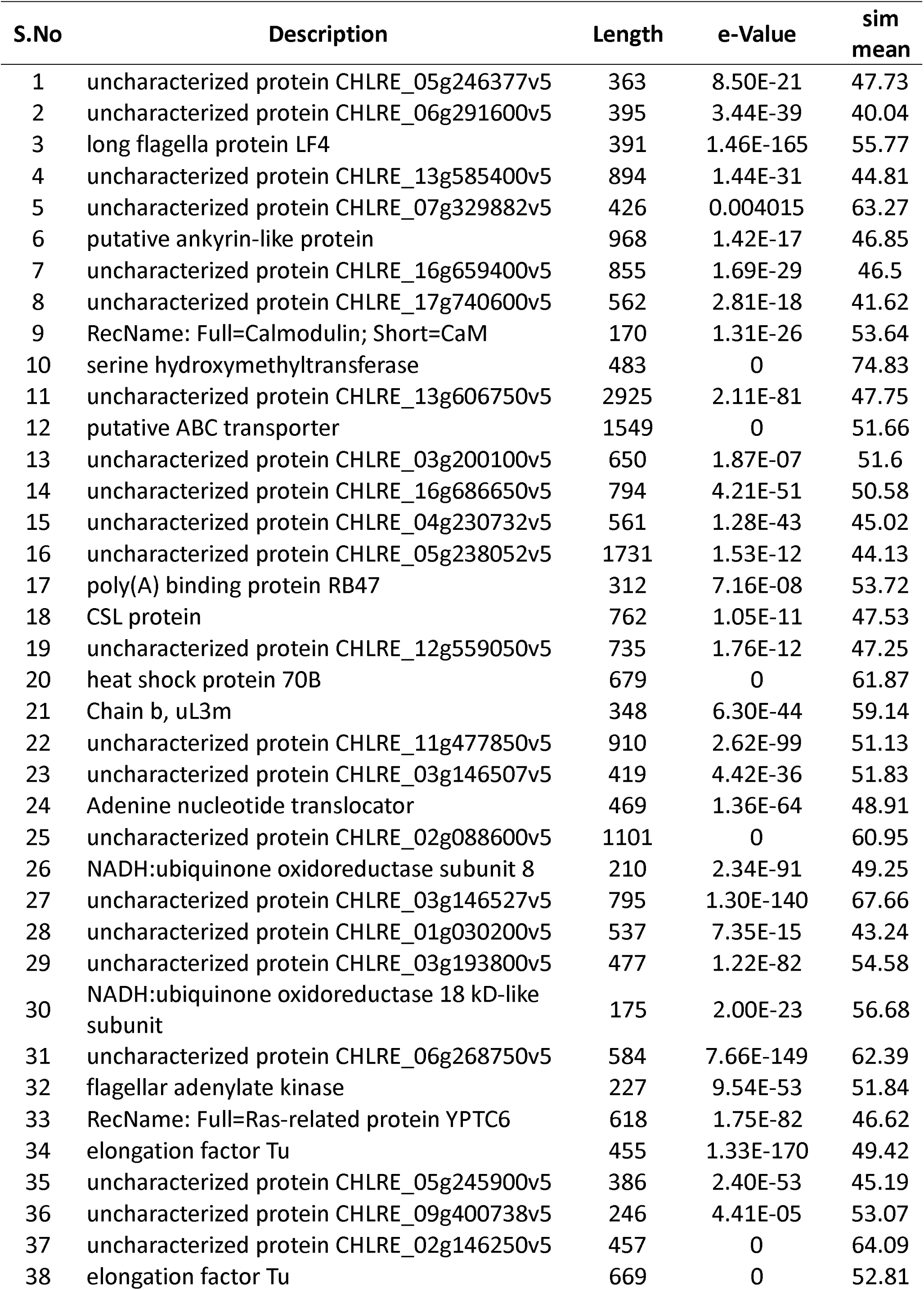
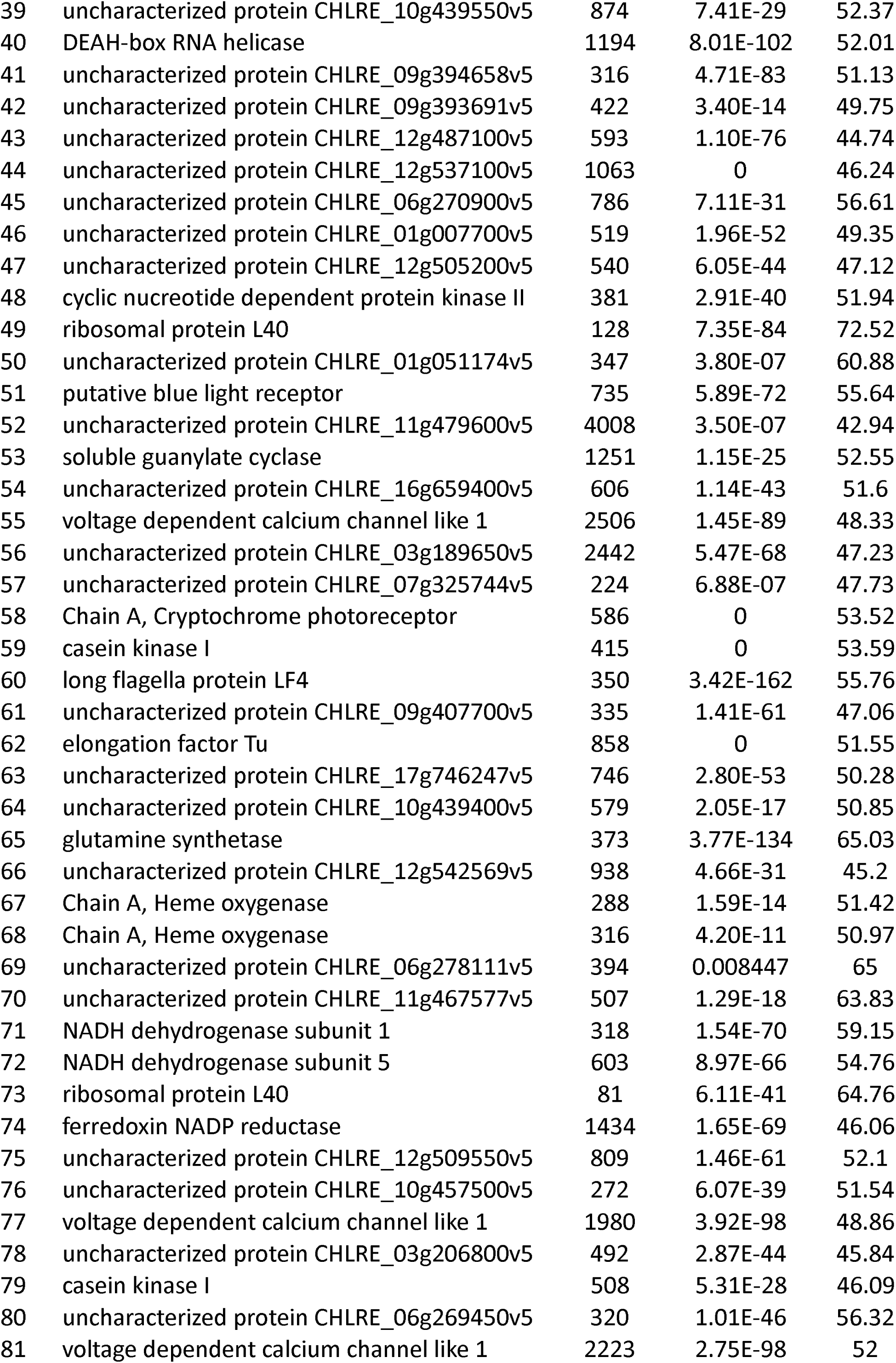
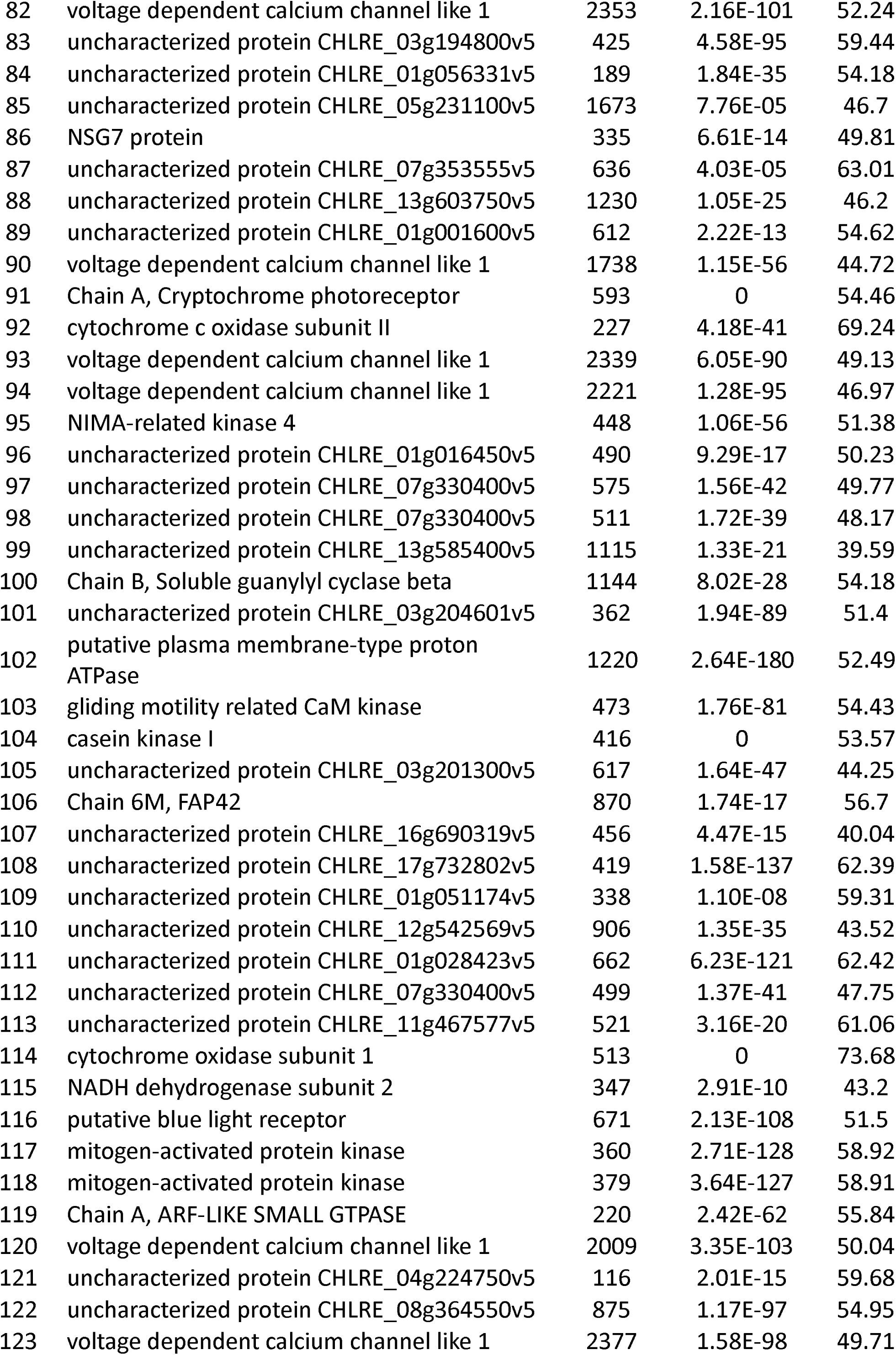
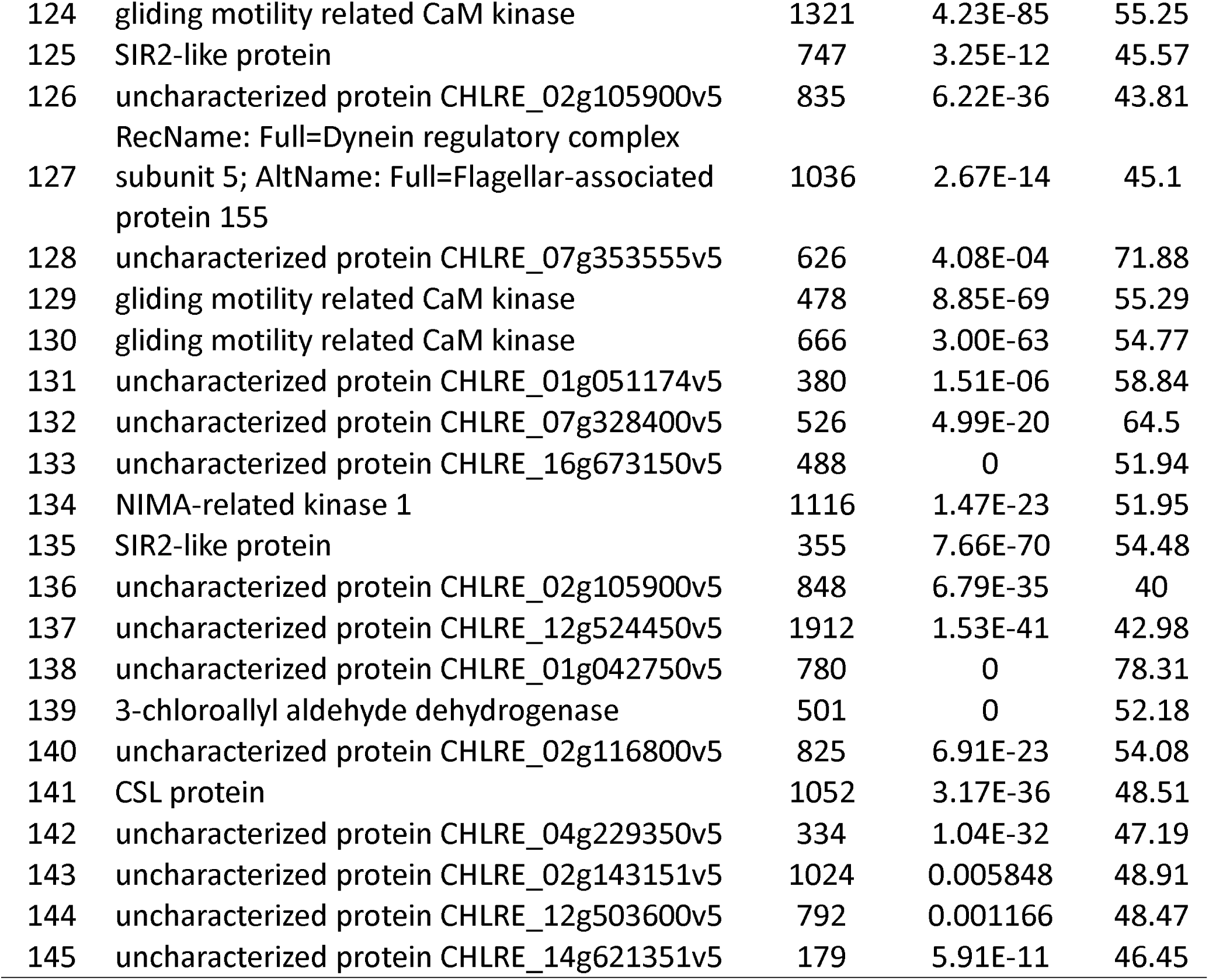
Hits obtained by performing sleep genes BLAST against *Chlamydomonas* genome.

This suggests a potential functional connection with other sleep genes that have already been identified. One of the several proteins that have yet to be fully characterized is CHLRE_07g325744v5, and this protein exhibits a similarity of 63.01%. This protein may have novel sleep-related activities that still need to be explored in Chlamydomonas. In addition to the uncharacterized proteins, several other proteins have already been characterized as elongation factor Tu (CHLRE_07g353555v5) and NADH.

Among the superfamily of enzymes are oxidoreductase (CHLRE_03g146527v5) and casein kinase I (CHLRE_01g016450v5). These proteins have been attributed to having defined roles in crucial cellular processes, and their inclusion in our analysis implies their potential involvement in sleep pathways. Anecdotal evidence shows that the casein kinase that controls circadian rhythms exhibits a sequence identity of 53.57% and an e-value of 0 (Zheng et al. 2014). This discovery demonstrates a strong correlation between the normal functioning of casein kinase I and sleep regulation in various organisms, augmenting the credibility of the identified findings. The table also indicates the presence of many associations between voltage-dependent calcium channels and adenylate kinase. These proteins participate in signal transduction and energy synthesis that are closely linked to sleep. The protein with the accession number CHLRE_01g028423v5, known as the voltage-dependent calcium channel-like protein, evinces a similarity of 54.18%. The lowest E-value observed was 1.84E-35, which indicates a substantial likelihood of its involvement in calcium signaling. This protein may be associated with sleep-related processes.

The BLAST analysis has identified a group of possible homologs of sleep-related genes in *C. reinhardtii*. A cursory analysis shows that while certain proteins are shared across many species, there are still a significant number of unidentified proteins that could offer a novel understanding of the mechanisms governing sleep in tiny plankton. Further investigation is necessary to establish the functions of these proteins in sleep, especially to uncover new molecular pathways that control this important physiological process. Figure 1 presents distinct information regarding the characteristics and overall distribution of the BLAST hits. While Figure 1A displays the analysis of BLAST hits based on the number of hits acquired per query sequence, most of the query sequences have yielded between 2 and 6 positive matches, with a few producing as many as 17 matches. This suggests that specific sleep-related genes are highly conserved with their numerous homologs or matches in *C. reinhardtii*. On the other hand, less conserved genes elicit a specialized function and have only fewer matches.

**Figure 1:**
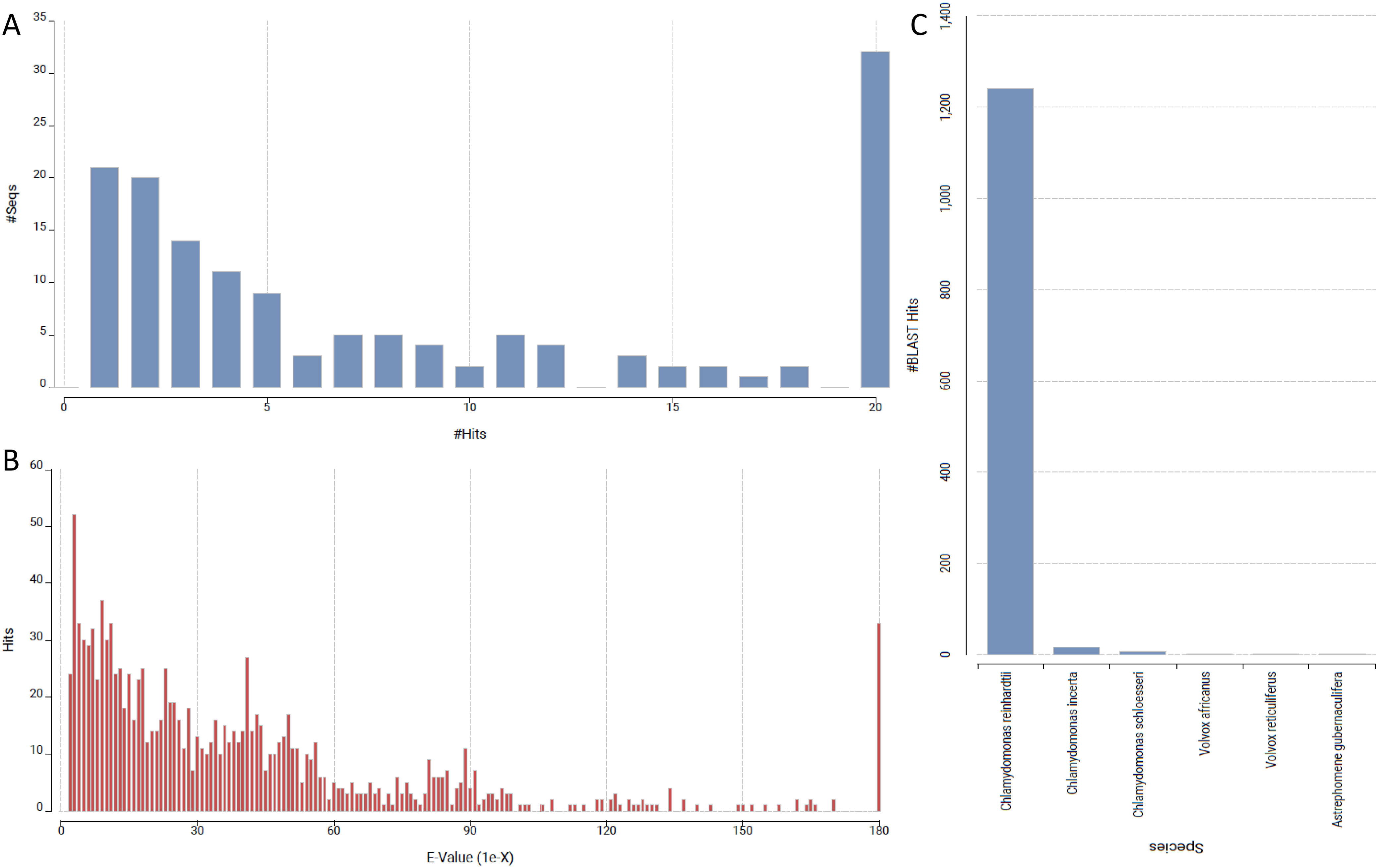
(A) Distribution of BLAST hit counts (#Hits) across query sequences. (B) Distribution of E-values for BLAST hits, indicating the significance of matches, with most hits having low E-values. (C) Species distribution of BLAST hits, with Chlamydomonas reinhardtii dominating the matches.

In Figure 1B, the histogram highlights the e-values detected inside the BLAST hits. Most hits exhibit a low e-value indicating a strong likelihood that the compared sequences could be homologs. Approximately 50% of the hits have an e-value close to zero, thus suggesting that they are substantially similar, and further suggesting that these hits are likely to be authentic homologs of the query sequences (Kerfeld and Scott 2011). However, a subset of findings with a higher e-value indicates that these sequences are less comparable and may represent more distantly related or potentially irrelevant sequences. The data shows that many alignments are moderately similar but not extremely divergent. Although there is a noticeable level of conservation in these sequences, there is still a significant amount of divergence that can be attributed to the various roles of these proteins. As observed in Figure 1C, BLAST hits throw light on the taxonomic distribution of the species. As anticipated, most hits are attributed to *C. reinhardtii*, given that the analysis was explicitly focused on this species. There are fewer instances of genetic matches from closely related species such as *Chlamydomonas incerta* and *Chlamydomonas stigma* and even fewer matches from more distantly related species like Volvox africanus and *Asterochloris glomerata*. These findings suggest that while the *C. reinhardtii* sleep-related genes have similar counterparts in closely related species, the level of conservation decreases as we look at species in more distant groups.

Figure 1 provides a comprehensive overview of the methodology employed for BLAST analysis and the distribution of similar sequences about the number of matches, expected value, percentage of similarity, and taxonomic classification at the genus and species level. The data demonstrate a moderate degree of conservation of sleep-related genes in *C. reinhardtii* and its relatives. These findings point to the need for further investigation of the functions of sleep genes in various species, notwithstanding the physiomorphic identity and taxonomic segregation.

### Distribution of sleep gene hits based on domain and families

Figure 2 describes the sequences based on the InterPro domains and families retrieved from the dataset. The content is partitioned into two panels: Panel A focuses on the InterPro (IPR) domains, whereas Panel B displays the InterPro families and the corresponding number of sequences (#Seqs) in each family.

**Figure 2:**
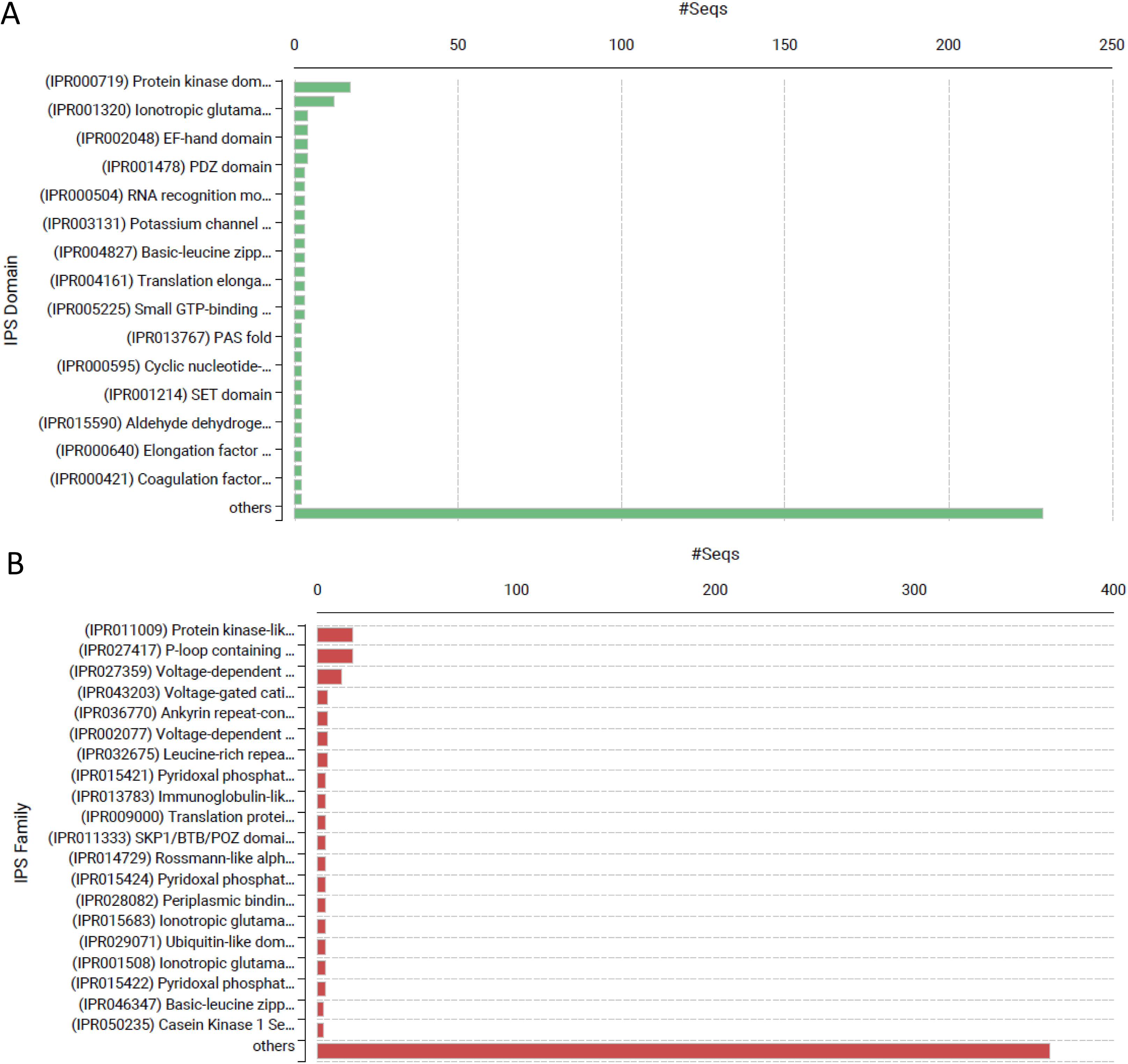
(A) Distribution of protein sequences across InterPro domains, with the majority belonging to the “others” category, followed by protein kinase domains and ionotropic glutamate receptor domains. (B) Distribution of protein sequences across InterPro families, showing the dominance of casein kinase I family members, with other families contributing fewer sequences.

Figure 2A displays the most representative InterPro domains that were acquired. The most prominent is the “Protein kinase domain” (IPR000719), exhibiting many associated sequences that suggest a significant amount of kinase activity within the dataset. ‘Ionotropic glutamate receptor domain’ (IPR001320) and the ‘EF-hand domain’ (IPR002048) observed here indicate a significant presence of ion signaling and calcium-binding proteins in the analyzed sequences. Additional domains present in the dataset are the ‘PDZ domain’ (IPR001478) and the ‘RNA recognition motif’ (IPR000504). However, they are rare. The presence of several domains relating to protein-protein interaction, signaling, and regulation points to an underlying functional diversity within the dataset. The ‘others’ group is broad and encompasses numerous under-represented domains, highlighting that there could be a wider diversity in domain types. As seen in Figure 2B, InterPro families are utilized to categorize sequences. The ’Protein kinase-like family’ (IPR011009) remains the most prevalent category and one that immediately draws attention. Following these families are the ’P-loop containing nucleoside triphosphate hydrolases’ (IPR027417) and ‘’Voltage-dependent calcium channel’ (IPR025523), which play a crucial role in cellular signal transduction and regulation. Other well-known domains include the ‘Ankyrin-repeat-containing domain’ (IPR002110), which is involved in protein-protein interactions and signal transduction pathways, and the ‘Leucine-rich repeat’ (IPR001611), implicated with the protein-protein interactions. As in Panel A, the ‘others’ group in Panel B displays a multitude of protein families, albeit with lower frequency. This indicates that the dataset encompasses a wide range of protein families.

Figure 3 illustrates the widespread presence of kinase activity, protein interaction domains, and protein families. It highlights the abundance of these proteins and showcases their functional versatility and richness within the dataset. The prevalence of sequences associated with kinases and signaling-related families indicates that in the researched system there is no dearth in signaling pathways and regulatory activities as well. This distribution provides insight into the potential biological tasks and pathways where these proteins may intercede to facilitate functional characterization.

**Figure 3:**
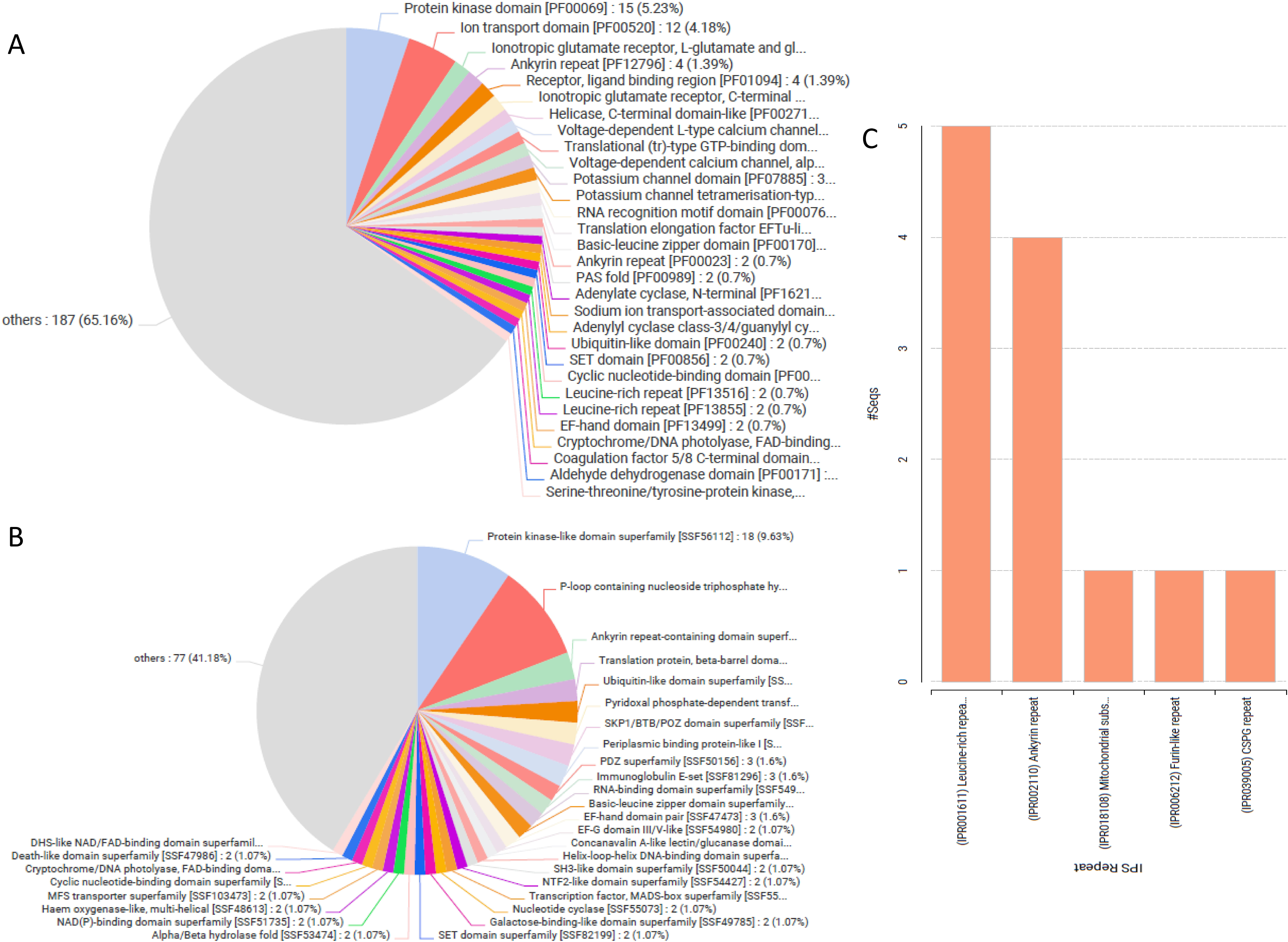
(A) Pie chart showing the distribution of protein sequences across Pfam domains, with the largest proportion categorized as “others,” followed by protein kinase, ion transport, and ionotropic glutamate receptor domains. (B) Pie chart displaying the distribution of protein sequences across superfamilies, with the largest group classified as “others,” followed by protein kinase-like, P-loop containing nucleoside triphosphate hydrolase, and ankyrin repeat superfamilies. (C) Bar chart of InterPro repeat sequences, showing the most common repeat types, with leucine-rich repeats being the most abundant.

Figure 3A-3C illustrates the allocation of protein domains and families within the dataset using pie charts and bar graphs. The analysis is segmented into three panels: the protein domain distribution, the protein family distribution, and the domain family distribution.

Figure 3A features a pie chart that illustrates the sequences analyzed about the PFAM protein domains. The most frequently encountered domain is the “Protein kinase domain” (PF00069), which encompasses five proteins. 23% of the total sequences indicate the significance of kinase activity in the analyzed input set. The presence of other domains, such as the “Ion transport domain” (PF00520) and “Ionotropic glutamate receptor” (PF00060), further emphasizes the importance of ion transport and signaling in this group of proteins. Nevertheless, most sequences (65.16%) are classified under the “others” category, which includes the less common domains in the sample, indicating a significant level of variety.

FIgure 3B displays the identical pie chart but specifically focuses on the PFAM protein families to which the sequences belong. The “Protein kinase-like domain superfamily” (IPR011009) is the most prevalent family on the list, with a count of 8. The findings from Figure 3A support the importance of kinase-related functions. Additionally, the “P-loop containing nucleoside triphosphate hydrolase superfamily” (IPR027417) and “Ankyrin-repeat-containing domain superfamily” (IPR002110) are also found in higher proportions, indicating their involvement in nucleotide binding and protein-protein interactions, respectively. Like Panel A, most of the sequences (41.18%) belong to the “others” category, indicating a significant variety of protein families in the sample.

Figure 3C transitions to a bar graph format and examines specific recurring sequences identified in the provided dataset. The most commonly occurring repeats are “Leucine-rich repeat” (IPR001611), “Ankyrin repeat” (IPR002110), and “Mitochondrial substrate carrier family” (IPR018108). These repetitions are crucial in facilitating protein-protein interactions and several cellular transport mechanisms. Figure 3C also illustrates the presence of “TGF-beta” and “GS repeats,” indicating distinct functional contributions from these proteins.

Overall, Figure 3 emphasizes the ’practical specialization’ of the protein sequences in the dataset concerning kinase-related activities, ion transport, and protein-protein interactions. Consequently, the proteins examined have the potential to be targets for in-depth analysis of the functional and structural aspects of biological processes and pathways related to the formation of the investigated domains and families. Additionally, the prevalence of specific domains and families implies biological functions that may play a crucial role in the processes above.

The categories with the highest abundance are “Monoatomic ion transport” and “Transmembrane transport”. Cell division and Long-term memory also contribute smaller amounts (Figure 4A). Additionally, there are other processes related to cell regulation, such as “Positive regulation of canonical Wnt signaling pathway.” The remaining terms are classified as “others.”

**Figure 1:**
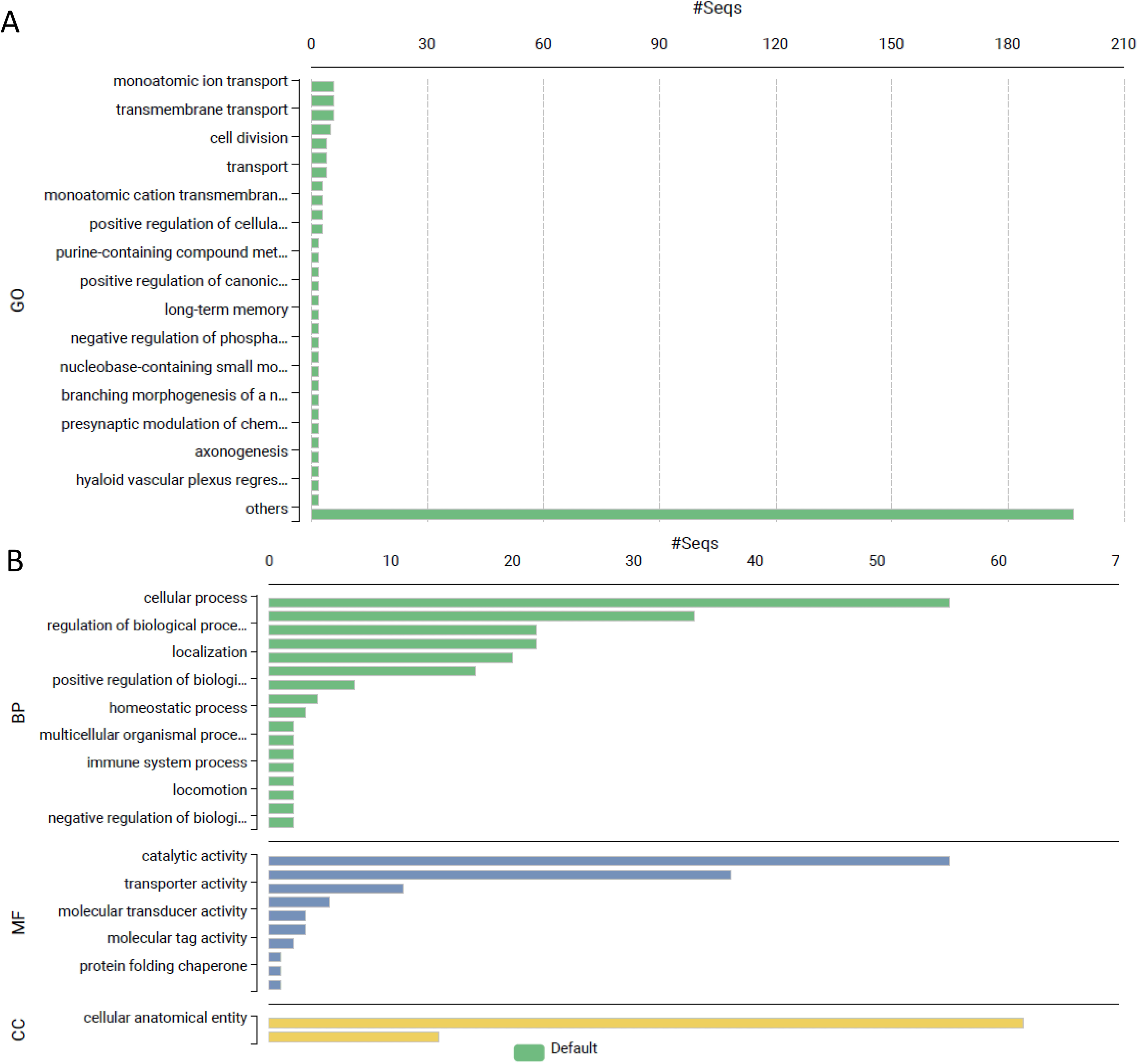
Gene Ontology (GO) annotation of the identified sequences. (A) Bar chart representing GO categories associated with molecular functions (MF), cellular components (CC), and biological processes (BP). The chart shows the number of sequences (#Seqs) assigned to each GO term. (B) Breakdown of the sequences within three primary GO categories: Biological Process (BP), Molecular Function (MF), and Cellular Component (CC). Biological processes such as “cellular process” and “regulation of biological process” are the most enriched, followed by molecular functions like “catalytic activity” and “transporter activity.” The “cellular anatomical entity” term dominates the Cellular Component category.

Furthermore, Figure 4B classifies GO annotations based on biological processes, molecular functions, and cellular components. The two most identified cellular processes in the BP category are controlling biological processes and localization. The MF category has a high degree of catalytic and transporter activity enrichment. “Cellular anatomical entity” is the most common term in the cellular component category (CC). The GO annotations are categorized and color-coded to indicate that these genes are inherently associated with various cellular and molecular biology aspects. According to this analysis, the genes examined involve several essential biological processes, molecular functions, and cellular components, highlighting the genes’ adaptability inside the cell.

### Pathway analysis

Figure 5 displays and reveals the number of paths in different categories in panels A and B. Panel A shows a comprehensive categorization of pathways into human diseases, organismal systems, and metabolism based on broad biological classification. Only “Human Diseases” and “Organismal Systems” have more than 60 paths among all the categories. Categories such as “Metabolism” are also significantly enriched, and the smaller groups like “Environmental Information Processing,” “Cellular Processes,” and “Genetic Information Processing” also exhibit many associated diseases. This indicates that most pathways are crucial for disease processes, organismal function, and metabolism.

**Figure 1:**
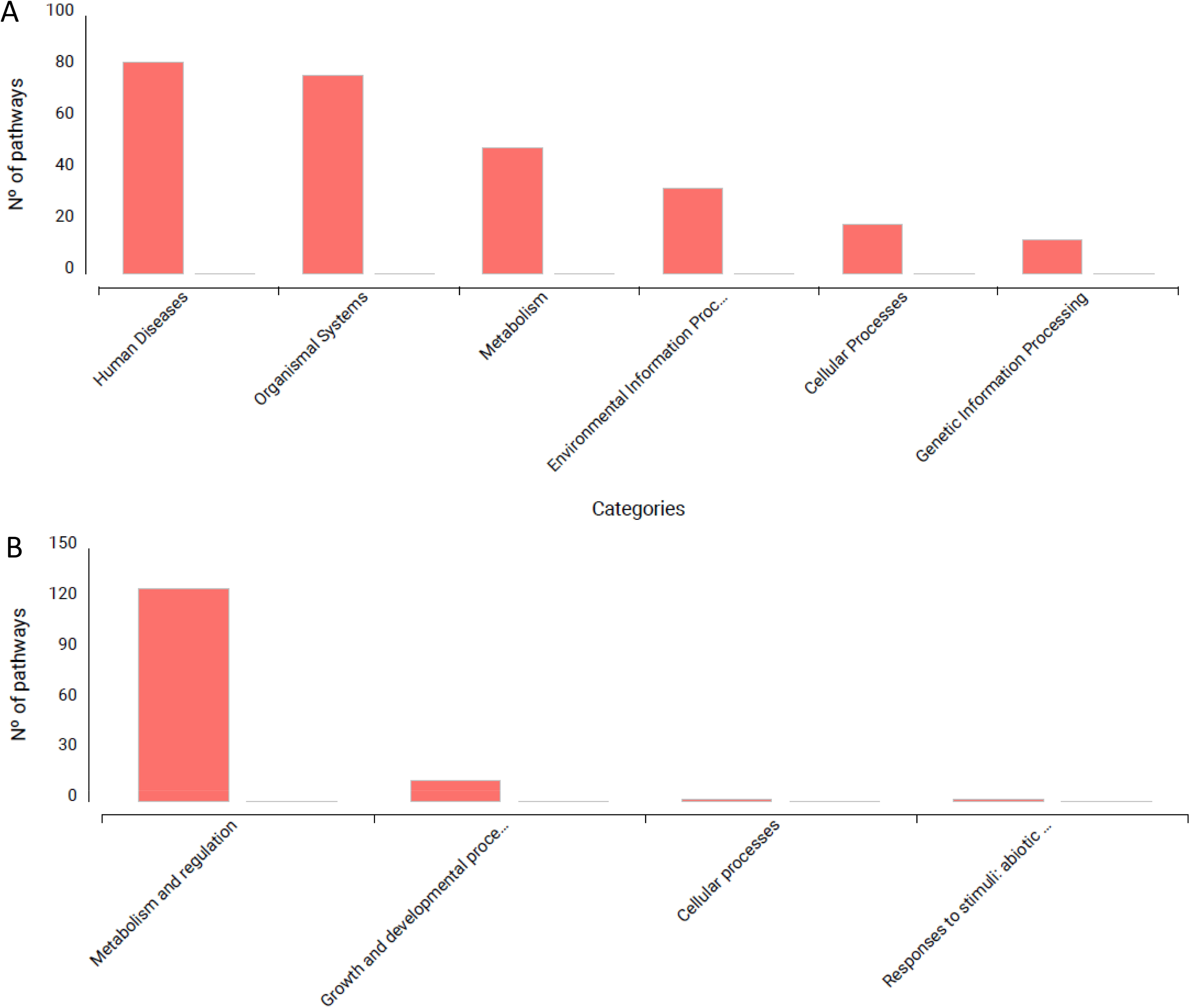
KEGG pathway classification of identified sequences. (A) Bar chart representing the number of pathways associated with different KEGG categories, including “Human Diseases,” “Organismal Systems,” “Metabolism,” and others. (B) Breakdown of pathways showing enrichment in “Metabolism and regulation” as the most dominant category, followed by minor contributions from categories such as “Growth and developmental processes,” “Cellular processes,” and “Responses to stimuli.”

Panel B enhances the inquiry by contrasting pathways associated with specific biological classifications such as “Metabolism and Regulation,” “Growth and Developmental Processes,” and “Cellular Processes.” The “Metabolism and Regulation” category stands out as the most prevalent, with more than 120 associated pathways. This indicates a significant presence of metabolic pathways. Additional categories include “Growth and Developmental Processes,” “Cellular Processes,” and “Responses to Stimuli.” However, other groupings, such as ’abiotic,’ are present but occur less frequently. This implies that the identified pathways have a high concentration of metabolic regulation. Furthermore, it elucidates that pathways primarily focus on metabolism and its regulation while recognizing their involvement in diseases, organism systems, and other biological processes. The analyzed data revealed a significant presence of the metabolism sequences category, which regulated numerous biological processes.

Figure 6 illustrates the distribution of sequences according to their functional categories, namely related to metabolism, cellular functions, and signaling pathways. This breakdown is presented in sub-figures A and B. Figure 6A displays three primary biological classifications: The three primary categories highlighted are “Metabolism,” “Cellular Processes and Signalling,” and “Information Storage and Processing.” The most dominant category is “Metabolism,” accounting for 35% of the sequences, indicating that the detected sequences are associated with metabolic processes. Following closely is the category of “Cellular Processes and Signalling,” accounting for 30% of the sequences that mark the presence of many cellular regulators. Approximately 25% of the sequences are categorized as “Information Storage and Processing.” These sequences feature genes that store, duplicate, and control genetic information. The dataset exhibits a noticeable bias towards metabolic and cellular metrics regarding the ’percent’ values.

**Figure 6:**
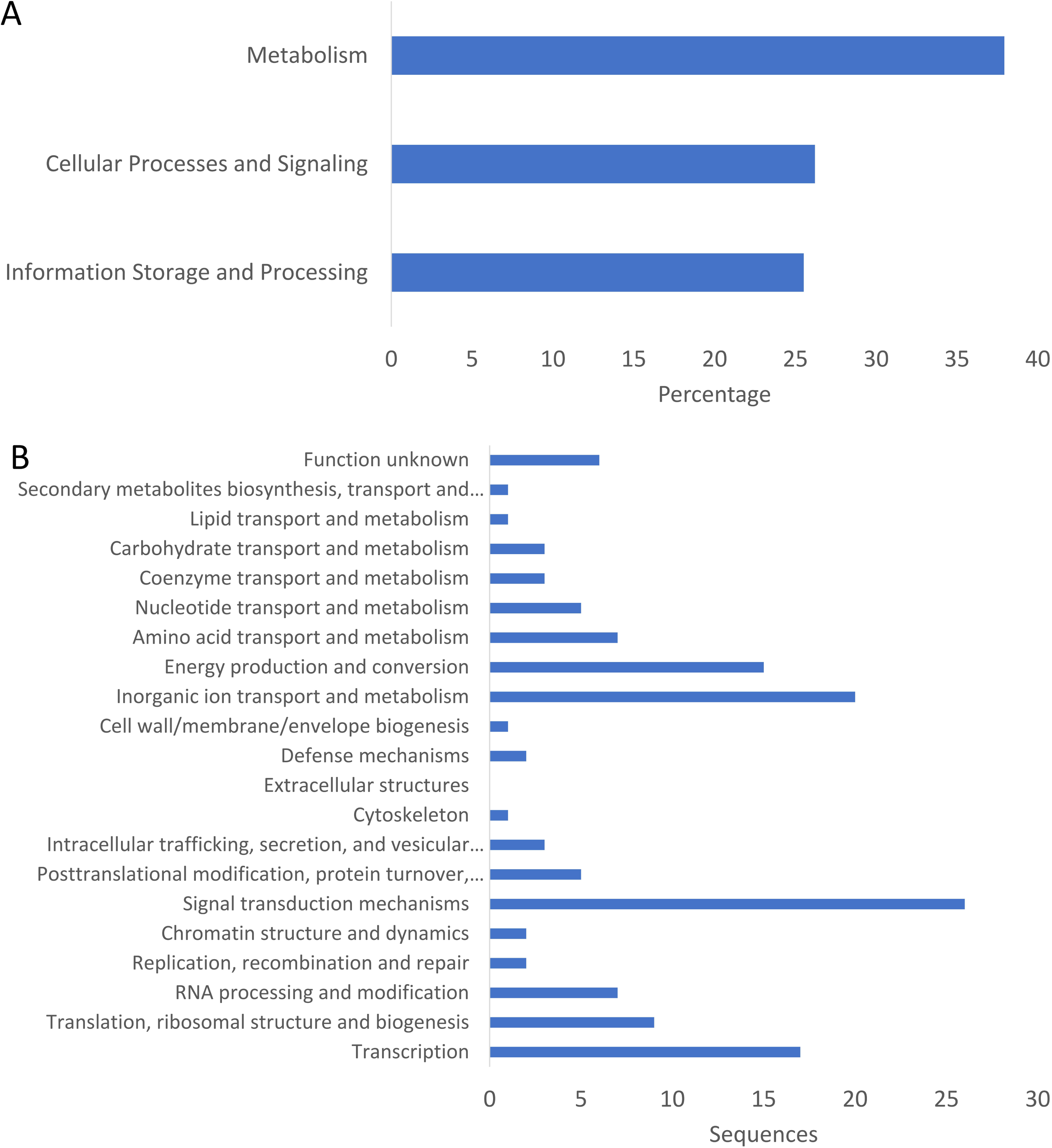
COG (Clusters of Orthologous Groups) functional classification of identified sequences. (A) Bar chart showing the percentage of sequences grouped into three major functional categories: “Metabolism,” “Cellular Processes and Signaling,” and “Information Storage and Processing.” (B) Detailed classification of sequences into specific COG functional groups, including “Energy production and conversion,” “Amino acid transport and metabolism,” “Signal transduction mechanisms,” and others. “Inorganic ion transport and metabolism” and “Signal transduction mechanisms” show the highest sequence counts.

Figure 6B provides a more comprehensive and extensive overview of sequences for each class. The main categories include “Energy Production and Conversion,” “Amino Acid Transport and Metabolism,” and “Inorganic Ion Transport and Metabolism,” each including around 15-25 sequences. Additionally, categories such as ’Nucleotide Transport and Metabolism’ and ’Carbohydrate Transport and Metabolism’ show significant upregulation, indicating a wide range of metabolic functions. The cellular functions of “Signal Transduction Mechanisms” and “Translation, Ribosomal Structure, and Biogenesis” are well represented with 20-30 sequences. Conversely, certain categories such as “Defence Mechanisms” and “Cytoskeleton” have fewer sequences, suggesting they are less critical in this dataset. Overall, the diagrams and illustrations show a significant emphasis on metabolism and energy provision with cells’ leading roles. Additionally, the dataset contains many representations of genetic information processing, signaling, and protein synthesis, indicating its diverse nature.

## Discussion

The results obtained from our BLAST analysis demonstrate that *C. reinhardtii* possesses many homologs similar to known genes associated with sleep, highlighting the evolutionary preservation of sleep pathways across various taxa. The detection of uncharacterized proteins, namely CHLRE_16g686650v5 and CHLRE_07g325744v5, exhibiting substantial e-values and similarity scores implies that these proteins might have essential but undiscovered functions in the regulation of sleep. The significant level of sequence similarity, especially in proteins that have yet to be well-studied, emphasizes the possibility of finding new and unique information inside an organism consisting of a single cell. Considering the substantial sequence alignment and e-values, a more thorough examination of the functional activities of these proteins could provide a novel understanding of the molecular mechanisms that govern sleep.

In the dataset, the presence of well-defined proteins, such as elongation factor Tu, NADH, and casein kinase I (CK1), provides evidence that sleep regulation encompasses a diverse range of conserved biological mechanisms. CK1, specifically, has been linked to the regulation of circadian rhythms, which are closely connected to sleep (Francisco and Virshup 2022). The high degree of similarity in the sequence and the extremely low e-value of this protein in *C. reinhardtii* strongly support its possible role in sleep-related pathways. This implies that similar regulatory mechanisms may be present even in single-celled organisms.

The existence of homologs for voltage-dependent calcium channels and adenylate kinase underscores the significance of signal transduction and energy production in sleep. The discovery of CHLRE_01g028423v5, a voltage-dependent calcium channel-like protein with a substantial similarity score, indicates that calcium signaling may be a common mechanism in different species for regulating sleep. This discovery is consistent with previous studies on Drosophila, which have demonstrated that calcium signaling impacts sleep patterns (Arnon et al. 1997).

Examining the domain and family distribution shows a notable frequency of kinase activity, ion transport, and protein-protein interaction domains, such as the protein kinase domain and the EF-hand domain. These findings indicate that signaling pathways and regulatory processes play a crucial role in the sleep mechanisms of *C. reinhardtii*. Numerous protein kinase-like families provide more evidence for the involvement of phosphorylation in regulating sleep-related activities, which is in line with similar discoveries in other animals.

Our examination of pathways highlights the significance of regulating metabolism and cellular activities in sleep. Identifying pathways linked to metabolism and regulation and the presence of signaling and cellular process pathways suggests a strong connection between sleep-related mechanisms in *C. reinhardtii* and the organism’s metabolic state. The presence of pathways associated with disease and organismal systems indicates that the proteins found may have further roles in broader biological activities, connecting sleep to overall cellular health and balance.

In summary, our study thoroughly examines possible counterparts of sleep-related genes in *C. reinhardtii*, emphasizing both shared and new aspects of sleep regulation. The notable existence of uncharacterized proteins (Table 1) presents promising prospects for more investigation, potentially revealing novel molecular pathways implicated in sleep. Preserving specific pathways and identifying distinct domains and families linked to sleep-related functions indicate that *C. reinhardtii* could be a valuable model for investigating the fundamental mechanisms of sleep. This research provides helpful insights that apply to more intricate organisms. Future study should prioritize the functional validation of these proteins and their involvement in sleep since this could result in a more profound comprehension of sleep’s evolutionary origins and biological importance.

Chlamydomonas has gained prominence as a model organism in biology and medicine; hence, after reviewing the study by Li et al. (Li et al. 2004), we aimed to determine whether Chlamydomonas possesses genes that are involved in sleep regulation. This research identifies 145 human gene orthologs and proteins characterized by distinct InterPro domains. The most significant share is attributed to the “others,” followed by protein kinase domains and ionotropic glutamate receptor domains, crucial for sleep signaling pathways and synaptic transmission associated with sleep. InterPro’s casein kinase I family members are disproportionately represented, indicating their essential role in regulating circadian rhythms and sleep-wake cycles. Other families contributed fewer sequences, yet these findings suggest that genes from Chlamydomonas may influence sleep patterns and offer distinct perspectives on sleep organization in simple and complex species.

Sleep is not a unitary state. The ability of animals to engage in unihemispheric (asymmetric form of NREM or SWS) sleep (Rattenborg et al. 2000, 2016; Mascetti 2016; Tamaki et al. 2016), a continuous activity in cetaceans after birth (Lyamin et al., 2005), the ability of the bird known as the Alpine swift to remain airborne for 200 days and to cover approximately 10000 kilometers in a non-stop flight (Tachymarptis melba) (Liechti et al. 2013) or post-flight recovery (Schwilch et al. 2002; Rattenborg et al. 2016) or for that matter, to somehow circumvent the necessity for sleep altogether e.g., non-sleeping piscine species (Kavanau 1998, 2008), challenges the notion that sleep is conserved or is a vital necessity for sustaining. Such findings point to the conclusion that not all animals might preferentially sleep as humans do and thus show that the overall phenomenon of evolution permits some degree of variability.

That vertebrate sleep might have occurred in the earliest jawed fishes that lived in interactive ecosystems at least 450 Mya (million years ago), further, that invertebrate sleep is far older (Lee Kavanau 2005) offers considerable latitude for researching the phenomenon. Undoubtedly, the recent progress in sleep genetics in simple model species deserves due recognition and calls for an enhanced understanding of sleep itself. Even though the inherent genetic manipulability of simplicity in these creatures inherently restricts their capacity to accurately represent all elements of sleep in mammals (Rattenborg et al. 2008), it should be agreed that the cross-referencing of data on genome structure among species or phylogenomics holds promise for developing critical perspectives on this biological act. Efforts in this area would thus extend the applicability of the Human Genome Project and would potentially yield a wealth of knowledge useful for improving animal breeding and health (Graves 1998).

The findings of this analysis support the suggestion that sleep has evolved as a key strategy of rejuvenation and thus is essential for the preservation of life. Should it be a mutational outcome of higher forms of land-based life or an inherent survival strategy in all life can be verified by a comparative analysis. This is the approach taken in the present investigation. Comparative genomics is an incredibly useful tool for both investigating and advancing our understanding of the human genome and for shedding light on the mechanisms and causes of evolution. The central theme of comparative genomics is that the sequences that remain conserved are probably restricted (similar) because of evolutionary constraints, providing a perspective to probe across the biological spectrum. A DNA sequence may nevertheless provide important information about its function even if it does not match that of other species.

This raises the interesting question of whether sleep studies could test new lineage-specific modifications where the sequence has not demonstrated conservation yet due to a lack of time. Sequence identity is not always synonymous with conservation; a sequence’s beneficial alteration may only affect two or three of the four bases, whereas an adverse alteration may be fatal and result in the death of the individual or species in question. Medical genetics is greatly impacted by the comparative genomics discoveries, and this influence is only predicted to grow in the future. Comparative genomics employs evolutionary theory to glean insights into the function of genomic DNA sequences (Hardison RC 2010). One can infer chromosomal rearrangements, duplications, and deletions, as well as the speeds at which different sequences have evolved, by comparing DNA and protein sequences within and between populations within a species.

The functional characteristics of the DNA can then be predicted using this evolutionary reconstruction. While regions that confer an adaptive benefit when altered are likely to have larger divergence among species, sequences required for common functions among the species under comparison are expected to change little throughout evolution. Additionally, sequence comparisons can aid in the prediction of the function of a specific functional region, such as coding for a protein or controlling the expression of a particular gene. Consequently, the concept of sleep can be understood within this framework, which posits that sleep represents one of the most profoundly evolutionary ingrained yet simultaneously genetically refined behaviors.

Comparative genomics, which has grown to be an essential tool in biological research, is the study of the variations and similarities in the genome structure and organization in other organisms. It is a very potent method that yields biological insights that are not possible with any other approach. Numerous genetic and physical methods can be used to determine the arrangements of genes and other DNA sequences, with resolutions ranging from the single base pair to the overall cytological level (Graves 1998).

One of the main motivating factors is the need for a much more thorough understanding of the evolution process, both locally (what distinguishes related species from one another) and globally (the emergence of the major groups of organisms). The requirement to translate DNA sequence information into a protein with a known function is the second motivator. The reasoning for this is that DNA sequences that encode crucial biological processes have a higher probability of being conserved across species compared to noncoding or redundant sequences. According to Bofelli et al. (2004)(Boffelli et al. 2004), conserved sequences with likely important functions can be found by comparing the genomes of extremely distantly related species, such as fish and mammals. Comparative genomic analysis has the potential to yield precise estimates of gene content and orthology, enhance comprehension of the evolution of genome architecture and the degree of genomic synteny (refers to the conservation of the order of genes on chromosomes between different species) among species, and identify the percentages of genomes evolving under evolutionary constraints (Craig et al. 2021).

Moreover, whole-genome alignments (WGAs) between various species can be constructed, and conserved elements (CEs) in noncoding regions can be found for several of the most extensively researched lineages. The idea that sleep either underpinned or made possible the process of how single cells evolved into sophisticated organisms is opposed by biologists who believe that sleep is a controlled phenomenon, and thus of some but not overriding importance. This is a debate that continues today, but one which will require further studies of additional genes and gene networks. It will take a lot more effort to prove that even significant evolutionary shifts can be achieved with relatively small genetic alterations.

## Conclusion

In summary, the human genome sequence and the technical and technological advancements that have sparked phylogenomics as a valuable tool for studying the evolution of vertebrates’ genomes will help pursue such queries. It could also help us understand the human genome study, which has the potential to help us understand human sleep, sleep disorders, and the inescapable relationship between the two. In both comparative genomics and phylogenomics one can use genetic data to infer relationships between organisms, as they are closely related fields. Conversely, phylogenomics expands on the discoveries made possible by comparative genomics to produce a more profound comprehension of the evolutionary past at the organismal and genomic levels. In this work, we have shown the advantages of phylogenomics in the context of sleep with the hope that the currently available information in comparative genomics could potentially address the issues of the study of sleep and sleep disorders in a new dimension.

Our phylogenomic analysis based on Chlamydomonas, has helped to elucidate the evolutionary aspects of sleep-related genes. Thus, by selecting 145 human gene orthologs in Chlamydomonas, we underlined important protein domains, such as protein kinase and ionotropic glutamate receptor domains, which play a fundamental role in sleep signaling pathways. Thus, our results indicate the evolutionarily related processes of calcium signaling and synaptic transmission involved in sleep regulation are conserved across phyla, and that simple model organisms like Chlamydomonas may be useful in elucidating higher-order, complex behaviors such as sleep. Further, by using phylogenomics, one can determine conserved domains involved in circadian rhythm and sleep-wake regulation, suggesting the evolutionary conservation of these mechanisms. Chlamydomonas do not possess the complexity of the sleep seen in vertebrates. However, knowing their genetic makeup can help researchers understand general molecular processes that may be used to understand sleep in more complex organisms. Several comparative and phylogenomic studies are promising in the context of human sleep disorders. They may open a new avenue for developing new therapeutic approaches based on the detection of the evolutionarily conserved genes and pathways. Thus, the use of Chlamydomonas and C. elegans in sleep research has remained a source of a rich system for understanding the genetic factors involved and the evolution of sleep.

## CRediT authorship contribution statement

SRP; SBC: Conceptualization and Methodology; SRP; KMS; SP: Data collection and review, KMS; SP: Software, SRP; KMS; GCA: Preparation of original draft; SP; SBC: Critical review and editing; SBC: supervision; Prior to submission, all authors (SRP; KMS; SP; GCA; SBC) have read and agreed to the submitted version of the manuscript.

## Author Agreement Statement

All authors affirm that the paper is their original work and that it has not previously been published nor is it presently being considered for publication elsewhere. Each author expressly declares that they contributed equally, read, evaluated, and accepted the final version of the paper, and agreed to be included as co-authors as per ICMJE guidelines. Each of the authors has approved the author sequence and the corresponding authors.

## Role of medical writer or editor

Not applicable.

## Ethical Statement

The study carried out by the authors did not involve the use of animals or human participants. Hence, no IRB approval was necessary for this work.

## Data availability

This study utilized the dataset that was originally prepared for our ongoing phylogenomics and bioinformatics investigation of sleep. Since this is a work in progress, no additional information will be available from the authors.

